# KG-Bench: Benchmarking Graph Neural Network Algorithms for Drug Repurposing

**DOI:** 10.1101/2025.10.13.682003

**Authors:** Siqi Wei, Christo Sasi, Jelle Piepenbrock, Martijn A. Huynen, Peter A.C. ’t Hoen

## Abstract

**Motivation:** Drug repurposing leverages existing drugs for new indications, accelerating drug development. Computational methods integrating diverse biological and chemical data can systematically prioritize repurposing candidates, but standardized benchmarks for deep learning evaluation are lacking. We present KG-Bench, a benchmarking framework for drug-disease association prediction using the Open Targets dataset. We constructed a knowledge graph (KG) of drugs, diseases, and targets, including annotations such as therapeutic area and molecular pathway, and ensured retrospective validation by leveraging regular dataset updates. To avoid data leakage, we removed redundant entities across splits.

**Results:** Benchmarking several graph neural network (GNN) architectures, TransformerConv achieved the highest performance (APR: 0.87). KG-Bench also assesses bias, node/feature importance, and uses GNNExplainer for interpretability. Our open-source framework enables fair, reproducible evaluation of graph-based drug repurposing algorithms.

**Availability and Implementation:** Data and codes are available at https://github.com/cmbi/Benchmark_GNN_OpenTargets.

## Introduction

Drug repurposing aims to identify new therapeutic uses for existing drugs by leveraging knowledge from previously known drug-target interactions[1]. Historically, successful drug repurposing projects have been conducted by biologists and pharmaceutical domain experts based on experimental evidence published in literature and clinical trial results[2, 3, 4, 5]. For rare and previously untreated diseases, insights from domain experts may nevertheless not be enough[6, 7].

Given the interconnected nature of drugs, diseases, targets, and biological pathways, deep learning provides a natural framework for drug repurposing by modeling complex biological relationships and predicting missing links that represent potential therapeutic opportunities. Deep learning methods allow automatic feature learning from massive datasets[8]. Furthermore, drug repurposing draws upon diverse data like chemical structures, genomic profiles, and protein-protein interactions. Given the diversity in the innate properties of the entities in biomedical datasets, deep learning frameworks learn from implicit complex relationships despite the variability in data source and data type[9]. The nonlinear modeling capabilities of deep learning architectures often result in superior predictive performance compared to traditional machine learning methods that rely on handcrafted features or linear models for identifying potential drug repurposing candidates[10].

Graphs can be used for the representation of functional, structural, and other complex information inherent in non-standard real-world data to train a GNN[11, 12]. A more enriched variant of a graph, which can be used for the representation of knowledge, is called a KG. The cumulative biological and pharmacological knowledge from clinical and molecular interaction studies can be extracted from databases such as ChEMBL[13], DrugBank[14], PubChem[15], and PDB[16]. KGs are composed of nodes, edges, and their features. Nodes are entities labelled with unique keys such as a ChEMBL ID for a molecule, an Experimental Factor Ontology (EFO) ID for a disease, and an Ensembl Gene ID for a gene. Edges are entities used to represent the relationship between two node entities. For example, if a molecule *m* is approved for the treatment of a disease *d*, it can be represented as (*m, d*). In addition, the properties or characteristics of the nodes and edges (features) are encoded as tensors that can capture and represent the various features. Feature types may include Boolean features, numerical features, categorical features, and text features. By encoding features in different ways, graphs can effectively represent complex entities in a format that models can process. This enables models to learn and distinguish the properties of different nodes, thereby implementing tasks such as classification, prediction, and reasoning in the graph structure[17]. Drug repurposing candidate selection based on biomedical KGs has gained popularity in recent years because they facilitate the representation of intricate biological relationships, such as drug–target interactions, disease–gene associations, shared molecular pathways, and phenotypic similarities between diseases[18, 19].

Graph learning algorithms can be trained on KGs to predict new links between previously unconnected nodes. These methods leverage the topology and features of a graph and iteratively propagate information from a node to its neighbors[20]. While the complexity of deep learning can hinder interpretation, GNNs offer reasoning opportunities. This contributes to the understanding of mechanisms of action and builds greater confidence in predictions[21]. GNNs are expected to make better predictions when the edge connectivity in a graph is dense, as it provides more information for the model to learn from. This is especially true in biomedical KGs where multiple relationships between nodes can offer valuable insights. For instance, when predicting drug–disease associations, GNNs can leverage not only direct drug-target interactions but also indirect pathways through shared protein targets, gene co-expression patterns, and disease phenotype similarities[22].

In a KG-based prediction of drug repurposing candidates, representations of drugs and diseases that are already linked are expected to aid the GNN in predicting additional links for these drugs[23]. The loss function penalizes the model for incorrectly predicting links that do not exist and for missing links that do exist in a validation or test set. A positive link prediction occurs when the model accurately predicts a link that exists in the validation dataset. Conversely, a negative link prediction is when the model correctly predicts the absence of a link that does not exist in the validation dataset. However, biomedical KGs typically contain only observed positive associations (known drug-disease relationships), while the vast majority of possible drug-disease pairs remain unobserved and unlabeled, and are typically sampled from the non-existent links. This creates an imbalanced learning scenario and affects the accuracy and generalization capacity of drug repurposing tasks[24, 25].

The Open Targets platform provides a single source of more than ~20 publicly available expert-annotated biomedical datasets. The dataset contains some salient features that can be useful in creating a graph representation of the data and a deep learning framework for drug repurposing. The Open Targets dataset is updated every three months with new annotations based on real-world discoveries of gene-disease associations and drug-disease approvals based on successful clinical trials reported in the literature. This allows users in principle to train models on historical data and validate and test the models using links added in subsequent updates. Such a train-validate-test framework is useful for drug repurposing predictions. An analysis of the Open Targets Dataset[26] showed that 33 out of 50 (66 %) drugs approved by the FDA (Food and Drug Administration) in 2021, had either one or all the following features: the genes encoding their primary assigned targets had previously been associated with the drug’s indication; proteins known to physically interact with the drug targets had established associations with the indication; some phenotypes were closely related to the drug indication and were genetically related to the drug target.

We have witnessed the development and publication of a large number of GNN-based algorithms for the prediction of drug repurposing candidates[10, 22, 27, 28, 29]. The performance of these algorithms is difficult to compare due to the absence of a gold standard dataset. The reported performance is often overestimated due to the absence of external validation sets, bias, and data leakage between training and validation datasets[30]. To address this gap, we developed a benchmarking framework that enables fair and realistic evaluation of GNN-based prediction tools. Our framework is built on KGs constructed from multiple time-stamped versions of the Open Targets dataset. This design allows models to be trained on historical data and validated or tested on future updates, simulating real-world scenarios where predictions are evaluated against newly reported drug-disease associations.

## Methods

### Framework Overview

A graphical representation of our benchmarking framework, named KG-Bench, is provided in Figure 1.

**Fig. 1.**
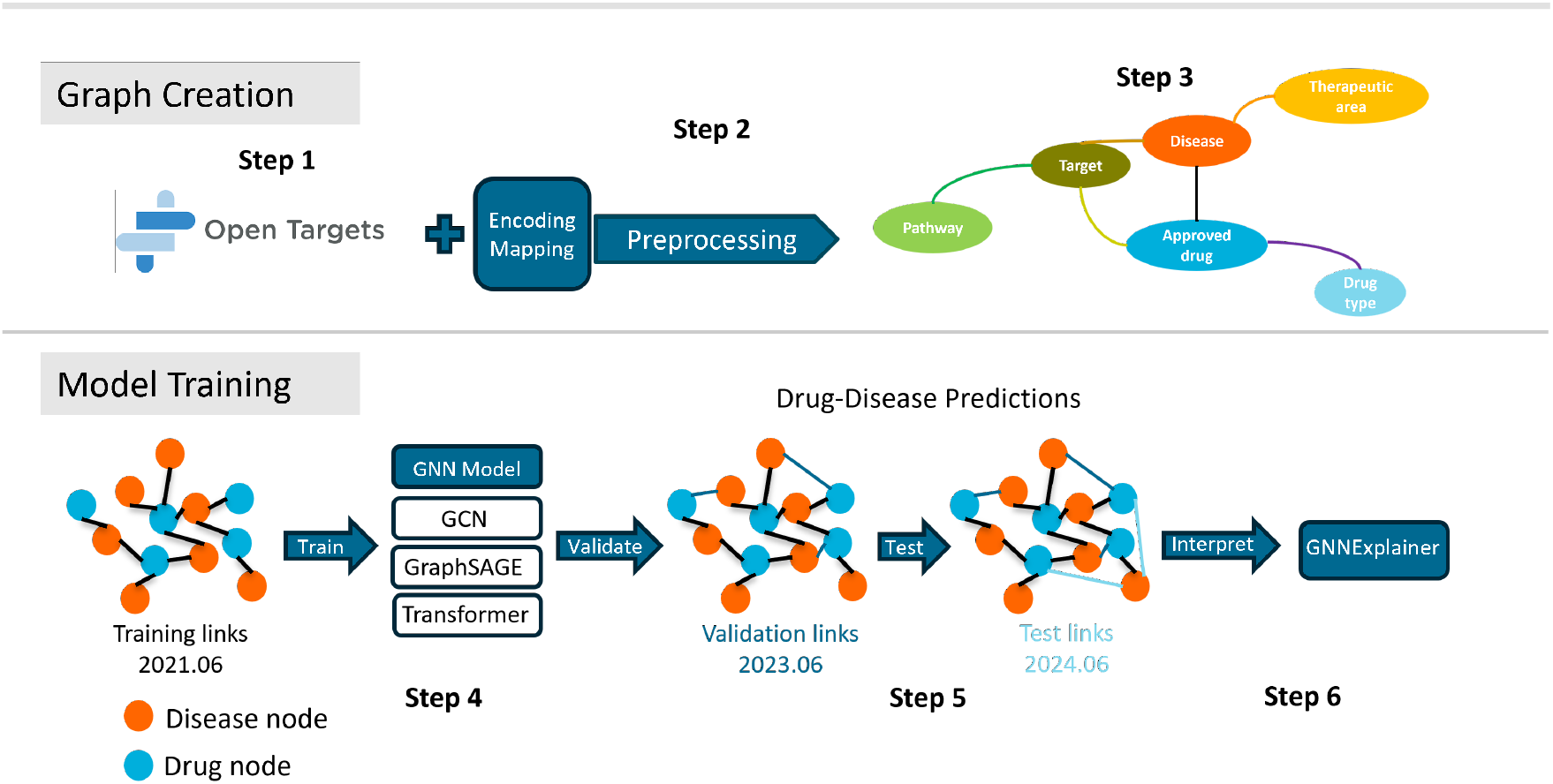
Benchmarking framework for evaluating GNN algorithms on drug repurposing using Open Targets KGs.

Step 1: The Open Targets dataset is queried to obtain the main entities and related annotations required to build the KG, followed by determining the nodes, features, and edges to extract from the graph, as described in Data Selection.

Step 2: The nodes, edges, and features are uniformly preprocessed and encoded. This involves cleaning the raw data, handling duplicates and missing values (see Data Preprocessing), normalizing column values based on their characteristics, and converting the results into tensor formats suitable for model input.

Step 3: The KG is constructed; the drug-disease edges in this version are the training set.

Step 4: We keep the nodes in the training set unchanged and focus on the drug-disease relationship edges added to these nodes in subsequent versions. The number of newly added edges in different versions is compared, and the versions with relatively large increments are selected as the source of validation and test sets. These newly added drug-disease edges are divided into validation and test sets respectively, and finally form a complete training, validation and test triplet together with the original training set.

Step 5: We use the KG from Step 3 to train multiple GNN models to effectively predict potential drug-disease links, and evaluate the prediction performance of the models on validation and test sets to test their generalization ability to new drug-disease relationships.

Step 6: We apply GNNExplainer to interpret the predictions by identifying key nodes and edges that influence drug-disease links. The results are statistically analyzed to assess the importance of different types of nodes and ensure the reliability of the explanation.

### Knowledge Graph

#### Data Selection

We first examined the parquet datasets (from: https://platform.opentargets.org/downloads) to identify relevant data for graph construction. Column names and data types were exported to CSV files and annotated as nodes, features, or edges based on requirements (see Data Preprocessing Section). We then established mappings between these entities and their attributes. The selection criteria are described below.

We established three types of core nodes: Disease, Drug and Target. In terms of drug representation, the platform adopts the molecular structure definitions used by ChEMBL, where a “parent molecule” refers to the original, unmodified form of the active ingredient, and a “child molecule” includes chemically modified variants such as salts or esters. To ensure comprehensive initial coverage of the drug space, both parent and child molecules were included. We selected only approved drug nodes, which are defined as compounds with a clinical trial max phase value of 4 for at least one indication. We further selected diseases and phenotypes that have at least one therapeutic relationship with approved drugs. In addition, we introduced three additional nodes, Drug Type, Therapeutic Area, and Pathway, to prevent some isolated triplets (Drug:Disease:Target) and increase connectivity in the KG. Among them, Drug Type is used to distinguish drug categories (for example, small molecules, antibodies, and oligonucleotides), Therapeutic Area is used to mark disease types (for example, infectious diseases, endocrine system diseases), and Pathway indicates the biological pathway in which the gene is located based on the Reactome platform.

The following undirected edge relationships were selected: drug-disease edges, which reflect treatment-related interactions; drug–target edges, representing the molecular targets of drugs; disease–target edges, which indicate known or predicted associations between diseases and biological targets like proteins; disease–therapeutic area edges, linking diseases to broader clinical areas; target–pathway edges, capturing the involvement of targets in biological pathways; and drug–drug type edges, specifying the classification of drugs (e.g., small molecule, antibody). The datasets and the columns selected for graph creation are listed in Table 1.

**Table 1.** Graph entities selected for input graph creation, split into node and edge types.

| Node Entities |  | Edge Entities |  |
| --- | --- | --- | --- |
| Dataset | Node Type | Dataset | Edge Type |
| Known drug | drug | Known drug | drug - drug type |
|  | drug type |  | drug - target |
| Target | gene | Target | target - pathway |
|  | pathway |  |  |
| Disease / Phenotype | disease | Disease / Phenotype | disease - therapeutic area |
|  | therapeutic area |  |  |
|  |  | Indication | drug - disease |
|  |  | Association score | disease - target |

Note: Node types include drugs (identified by ChEMBL compound IDs), drug types (identified by drug modality categories), genes or targets (identified by Ensembl gene IDs), pathways (identified by Reactome pathway IDs), diseases (mostly identified by EFO IDs, with additional cross-references to Orphanet, MONDO, and DOID IDs), and therapeutic areas (identified by EFO IDs, indicating therapy domains).

In the graph *G (N, E), N* is the number of nodes and *E* is the set of edges between nodes in the graph. Each node is represented by a one-hot encoded vector that represents its unique node type. The feature tensor of each node type is aligned to match the size of the feature tensor of the node type with the largest number of features. To reduce model complexity while preserving expressiveness, we selected a concise set of representative node features. The selected features of each node type are listed in Table 2. All existing edges between nodes are edges without a defined direction. This is achieved by including two sets of edges in *E* that represent interchangeable source node relationships. This allows our trained GNN to treat edges between nodes symmetrically, which is suitable for link prediction tasks.

**Table 2.** Dataset Feature Descriptions.

| Node | Column Name | Feature Encoding | Description |
| --- | --- | --- | --- |
| Drug | blackBoxWarning | boolean | Whether the drug has an FDA black box warning |
| Drug | yearOfFirstApproval | normalized | Year when the drug was first approved |
| Target | bioType | one-hot | Gene type |

#### Data Preprocessing

The Open Targets Platform provides its datasets in Apache Parquet format, which is optimized for efficient data storage and retrieval. The Python API for Apache Arrow, PyArrow v16.0.1, was used to generate queries for extracting data from the Parquet datasets[31].

To build a non-redundant drug-disease KG and predict whether there is an association between disease and drug, we performed multi-step cleaning and standardization operations on the drug and disease data provided by Open Targets. In the Open Targets platform, entities such as drugs, diseases, and targets are encoded using ontology IDs. Some IDs represent high-level entities, while others are more specific entities. Inconsistent IDs will cause the same entities in different hierarchies to be treated as multiple different nodes. This may not only cause the model to learn incorrect patterns, but also bring in an imperfect separation between training, test and validation sets. It may also lead to an overestimation of the performance of algorithms.

To ensure complete separation between training, validation and test sets of drug-disease relationships, we used the drug parentId only and removed all derivative drugs representing the same chemical entity. For example, sildenafil and sildenafil citrate share the same active compound, so keeping both would lead to data leakage where the model encounters essentially the same drug in both training and test sets, resulting in overestimation of performance. Diseases that lacked therapeutic field annotations were removed, and diseases related to the broad EFO term EFO 0001444, which represents measurement, an information entity, were excluded. We further filtered out IDs with prefixes UBERON, ZFA, CL, GO, FBbt, and FMA, which represent anatomical, developmental, or cellular entities. To prevent overestimation of the performance of models, we retained only the most specific level of disease from the disease ontology and removed all parent IDs (broader disease classifications). For all filtering steps, see Code availability, the graph creation script. The Open Targets database allows users to access datasets from July 1, 2019 to the present. The graph creation script accommodates changes in column names in the different versions of the Open Targets database.

#### GNN

The GNN models selected for benchmarking are:

- **GCNConv**: GCNConv is a semi-supervised learning algorithm for GNN based on a first-order approximation of spectral graph convolutions [32]. This model aggregates and normalizes the features of neighbors of a node to update its representation. It ensures that the updated representation of each node reflects both its features and those of its neighbors, scaled by their relative importance in the graph.
- **GraphSAGE**: The GraphSAGE operator is inspired by a graph isomorphism testing algorithm [33]. This model samples a subset of neighbors and aggregates their features, and the mean value is selected for the current task. This approach allows the model to generalize to unseen nodes by learning a function that generates embeddings based on sampled neighborhoods.
- **TransformerConv**: TransformerConv integrates GNNs with label propagation algorithms and introduces a masked label prediction strategy to prevent overfitting. This model uses attention mechanisms to focus on the most relevant neighboring nodes [34]. Attention mechanisms in

TransformerConv allow for a more nuanced aggregation of neighbor information compared with the other two algorithms.

All models produce 16-dimensional node embeddings. For link prediction, we use an inner-product edge decoder. For a candidate drug–disease pair (*u, v*) with embeddings *h*_*u*_, *h*_*v*_ ∈ ℝ^16^, we compute a scalar logit:

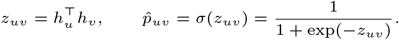

During training, we optimize the binary cross-entropy with logits:

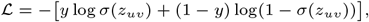

implemented with `BCEWithLogitsLoss` from PyTorch for numerical stability [35]. The loss is applied directly to the scalar logits *z*_*uv*_ (computed via inner product of 16-dimensional embeddings) for both positive and negative edges.

To ensure a fair and controlled comparison, all GNN models were trained using a unified hyperparameter configuration (Table 3). Specifically, we set the hidden feature dimension to 16 to reduce computational complexity while ensuring model expressiveness, thereby adapting to the data scale and training requirements of this task. The dropout rate is fixed at 0.5 to effectively suppress overfitting, while retaining sufficient model capacity. The number of convolutional layers is set to 2 to maintain a shallow graph structure, which is typically sufficient for link prediction tasks without long-distance information transmission [36]. In addition, all models apply layer normalization and ReLU activation functions to stabilize training and accelerate convergence. Early stopping patience, learning rate, data splits, loss function, and evaluation metrics were kept identical across architectures to isolate performance differences to model structure. Overall, these choices ensure a reasonable trade-off between model complexity, generalization ability, and reproducibility of results.

**Table 3.** Comparison of GCN, SAGE, and TransformerConv GNN architectures and edge decoder.

| Aspect | GCN | GraphSAGE | TransformerConv |
| --- | --- | --- | --- |
| GNN Layer Type | GCNConv | SAGEConv | TransformerConv |
| Layer Count | num_layers=2 |  |  |
| Hidden Dim. | hidden_channels=16 |  |  |
| Normalization | LayerNorm after each layer |  |  |
| Activation + Dropout | ReLU + Dropout (0.5) between layers |  |  |
| Multi-head Attention | – | – | heads=4, concat=False |
| Edge Decoder | Inner product ($h_u^\top h_v$) → scalar logit | | |
| Loss | BCEWithLogitsLoss on logits |  |  |

Note: All architectures share the same hidden dimension and optimization strategy; differences lie in the convolution operator and attention mechanism.

#### Ablation Studies

To evaluate whether specific connections between drugs and diseases affect the ability of the model to predict drug-disease associations, we shuffled the associations between drugs and diseases. In shuffling the edges between drugs and diseases, drugs were randomly reassigned to diseases while preserving both drug node degree (the number of diseases each drug is connected to) and disease node degree (the number of drugs each disease is connected to). The graph structure, negative training samples, validation and test set remained unchanged. To evaluate the contribution of node features, we removed all the node features or retained only the node type features.

### Negative Sampling Method

The Open Targets database contains only a set of positive drug:disease edges. To create a set of negative edges, we randomly sampled from the non-existent edges. We used balanced (ratio positive: negative = 1:1) and imbalanced (ratio positive: negative = 1:10 and 1:100) proportions for training, validation and test sets.

### GNNExplainer

We used GNNExplainer from PyTorch Geometric 2.7.0[37], a perturbation-based explanation method, to identify the importance of the nodes for the drug–disease prediction model[38]. This importance score quantifies how much each node (or node type) contributes to the prediction of the model. GNNExplainer optimizes learnable masks through gradient descent to identify minimal subgraphs that preserve original model predictions, thereby revealing mechanistically relevant graph components for each drug-disease pair. Comparative analysis of two neighborhood sizes was performed: 1-hop (immediate neighbors) and 2-hop (extended neighborhoods including pathway-mediated connections). For computational efficiency, we extracted k-hop subgraphs around target drug-disease pairs using subgraph sampling from PyTorch Geometric, with explanation generation performed on these reduced subgraphs before mapping results back to original node indices. Stratified random sampling was implemented at the confidence level to ensure unbiased representation. To quantify the variability and reliability of node type importance scores generated by GNNExplainer, we performed bootstrap confidence interval estimation. We generated 95% confidence intervals for node type importance rankings through bootstrap resampling (1000 iterations) of drug-disease pair explanations.

We applied a two-step validation framework to assess GNNExplainer performance. For baseline comparison, we assessed whether the model concentrates attribution on informative nodes rather than distributing importance uniformly. We generated random node-importance masks by sampling values independently on the same subgraph. We then used Mann-Whitney U tests to compare GNNExplainer’s average importance scores against this random baseline, verifying that explanations significantly differ from chance. For faithfulness testing, we determined whether top-attributed nodes are causally important for model predictions. We measured prediction performance drops after removing the top 20% of nodes by importance score. We compared these drops to those observed after removing randomly selected nodes using Wilcoxon signed-rank tests. The baseline comparison can verify that attributions differ significantly from random chance. Faithfulness testing can validate the predictive relevance of identified important nodes through systematic perturbation analysis.

## Results

### Retrospective Data Partitioning

As a training set, we selected the 2021.06 release of Open Targets, with 996 existing drug-disease edges after the strict data preprocessing criteria. Based on Figure 2, we selected the 2023.06 release as a validation set, with just over 400 newly added drug-disease combinations. This version is more than two years newer than the training set and is in the middle of the dataset timeline, serving as a suitable intermediate check to assess the generalizability with increasing data availability. As a test set, we used version 2024.06. This version helps assess the prediction or generalization ability of the model in the latest real-world data, reflecting the usefulness of the GNN prediction models in future real-world scenarios.

**Fig. 2.**
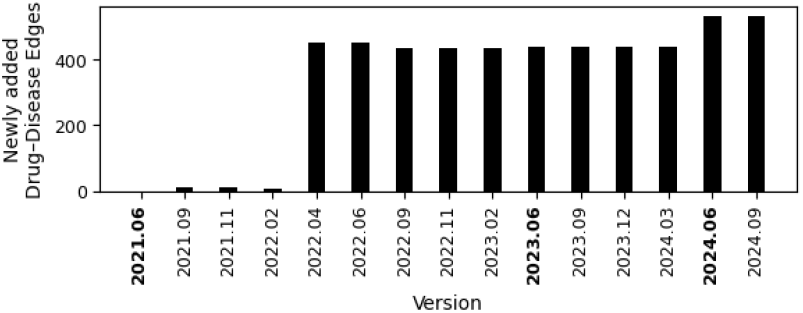
Newly added approved drug–disease associations since June 2021 across different versions of the Open Targets database

### Graph Statistics

The Open Targets KG used for training comprises 13,004 nodes and 106,793 edges (Table 4). The graph exhibits a scale-free topology, characterized by a small number of highly connected hub nodes and a large number of sparsely connected nodes. This is reflected in the average degree of 16.42 with a high standard deviation (76.93), indicating substantial heterogeneity in node connectivity (Figure S1). The negative degree assortativity (−0.133) suggests that hubs preferentially connect to low-degree nodes, reinforcing their role as bridges across otherwise weakly connected regions. The average clustering coefficient (0.011) is very low, consistent with the sparse modularity typical of biological networks. Hub drugs and diseases are highly connected, which will accumulate richer contextual information during the message passing process of the GNN, resulting in better learned embeddings and higher prediction confidence.

**Table 4.** Summary of node and edge types in the KG, including the number of instances and feature dimension.

| Node Types |  |  | Edge Types |  |
| --- | --- | --- | --- | --- |
| Node Type | Count | Feature Dim. | Edge Type | Count |
| Drug | 1926 | 3 | Drug - Drug Type | 1926 |
| Drug Type | 9 | 1 | Drug - Disease | 996 |
| Target | 726 | 2 | Drug - Target | 3192 |
| Reactome Pathway | 983 | 1 | Gene - Reactome Pathway | 3901 |
| Disease | 9336 | 1 | Disease - Therapeutic Area | 23801 |
| Therapeutic Area | 24 | 1 | Disease - Target | 72977 |

### Benchmarking GNN Algorithms

We used our benchmark KG to compare the performance of three GNN models that are commonly used in drug repurposing candidate predictions: GCNConv, TransformerConv, and SAGEConv. GCNConv is used as the baseline and we compare its performance against GraphSAGE and TransformerConv to evaluate the benefits of neighborhood sampling and attention-based message passing.

To simulate real-world scenarios, where the number of drugs that do not treat a disease greatly exceeds the number of drugs indicated for a disease, we validated and tested with multiple negative proportions. The negatives were derived from a random selection of edges that were not present in the training graph. It should be noted that a small fraction of these negatives may represent effective drug:disease combinations that have not been discovered yet. Along these lines, the false positives (FPs) are the most interesting fraction for follow-up research as they may represent new drug repurposing candidates. Figure 3 shows the Receiver Operator Curves (ROC) and Precision-Recall (PR) curves of the GCN, GraphSAGE, and TransformerConv models.

**Fig. 3.**
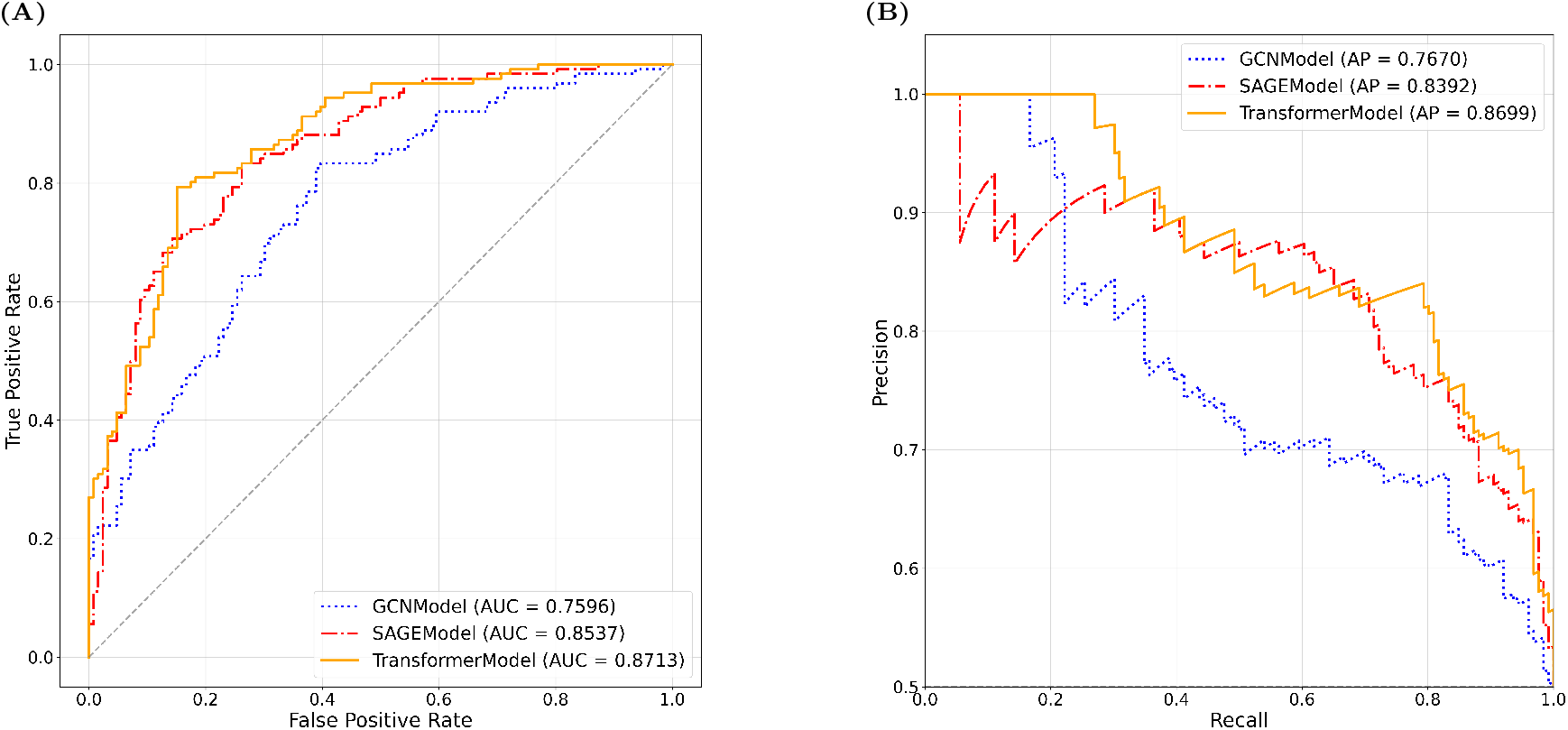
Model performance on test set evaluated with (A) ROC and (B) PR curves. GCN shown as dotted blue, SAGE as dash-dot red, Transformer as solid orange.

Overall, TransformerConv performed slightly better on all performance measures than GraphSAGE, which in turn performed better than GCN (Table 5). As expected, APR (area under the precision-recall curve) decreases as class imbalance increases because even a small fraction of FPs among a high number of negatives can significantly reduce precision, resulting in a lower APR. Recall is stable and high, indicating that the model can find most positive samples even in difficult, imbalanced environments. In summary, the TransformerConv model performs well in scenarios that reflect real-world scenarios.

**Table 5.** Model Performance on Test Set in Scenarios with Different Positive : Negative Ratios, with 95% Confidence Intervals.

| Ratio | Model | Sensitivity | Specificity | Precision | F1 Score | AUC | APR |
| --- | --- | --- | --- | --- | --- | --- | --- |
| 1:1 | GCN | 0.56 [0.47–0.65] | 0.75 [0.68–0.82] | 0.70 [0.60–0.78] | 0.62 [0.55–0.69] | 0.76 [0.70–0.82] | 0.77 [0.70–0.83] |
|  | GraphSAGE | **0.75** [0.68–0.83] | 0.77 [0.70–0.84] | 0.77 [0.69–0.84] | **0.76** [0.70–0.81] | 0.85 [0.81–0.90] | 0.84 [0.77–0.90] |
|  | TransformerConv | 0.66 [0.57–0.74] | **0.87** [0.82–0.93] | **0.84** [0.76–0.91] | 0.74 [0.67–0.80] | **0.87** [0.83–0.91] | **0.87** [0.82–0.92] |
| 1:10 | GCN | 0.56 [0.48–0.64] | 0.86 [0.84–0.87] | 0.28 [0.23–0.33] | 0.38 [0.31–0.43] | 0.81 [0.77–0.84] | 0.41 [0.33–0.50] |
|  | GraphSAGE | **0.75** [0.68–0.83] | 0.80 [0.77–0.82] | 0.27 [0.22–0.32] | 0.40 [0.34–0.45] | 0.87 [0.83–0.89] | 0.45 [0.37–0.54] |
|  | TransformerConv | 0.66 [0.58–0.74] | **0.91** [0.89–0.92] | **0.41** [0.35–0.48] | **0.51** [0.44–0.57] | **0.89** [0.86–0.92] | **0.59** [0.51–0.67] |
| 1:100 | GCN | 0.56 [0.48–0.65] | 0.87 [0.87–0.88] | 0.04 [0.03–0.05] | 0.08 [0.06–0.09] | 0.82 [0.78–0.86] | 0.11 [0.07–0.17] |
|  | GraphSAGE | **0.75** [0.67–0.83] | 0.81 [0.81–0.82] | 0.04 [0.03–0.05] | 0.07 [0.06–0.09] | 0.87 [0.84–0.90] | 0.11 [0.07–0.17] |
|  | TransformerConv | 0.66 [0.58–0.74] | **0.92** [0.92–0.93] | **0.08** [0.06–0.09] | **0.14** [0.11–0.16] | **0.90** [0.87–0.93] | **0.23** [0.16–0.31] |

**Note:** Values in brackets represent 95% confidence intervals. Best values for each ratio are highlighted in bold.

When interpreting the performance measures from drug repurposing prediction algorithms, FPs should not be considered as pure errors, but rather represent potential novel drug-disease associations. Analysis of FP predictions revealed significant consensus among the drug repurposing candidates predicted with the different model architectures (Table S1).

### Ablation Studies

We performed a set of ablation studies to evaluate the contribution of features, graph structure, and node-type information on the model (Table 6). The original model makes full use of the structure and multi-node characteristics. The full input setting (all features + true links) generally provides the best performance across models, representing the effective upper bound for this task. When the edges between disease and drugs are shuffled, the performance of all models drops to a level not better than random, indicating that graph structure is crucial for prediction and that prediction based on node features alone is not enough. GCN can still make accurate predictions using graph structure only, without the inclusion of node features. TransformerConv and GraphSAGE rely more on node features than GCN. With the node-type feature, the performance improves compared with the non-feature setting, indicating that mostly the node-type features, but not other node features, contribute to the prediction.

**Table 6.** Ablation study results showing the impact of features and edge topology on drug–disease prediction across different GNN models Setting TransformerConv GraphSAGE GCNConv.

| Setting | TransformerConv |  | GraphSAGE |  | GCNConv |  |
| --- | --- | --- | --- | --- | --- | --- |
|  | AUC | APR | AUC | APR | AUC | APR |
| All feat. + true links | 0.87 | 0.87 | 0.85 | 0.84 | 0.76 | 0.77 |
| All feat. + shuffled links | 0.52 | 0.52 | 0.44 | 0.49 | 0.47 | 0.48 |
| No feat. + true links | 0.48 | 0.50 | 0.52 | 0.51 | 0.82 | 0.78 |
| Node type feat. + true links | 0.73 | 0.71 | 0.86 | 0.86 | 0.80 | 0.79 |

In the remainder of the paper, we focus on TransformerConv, which achieved the highest performance in our benchmarks.

### Potential Biases

Model predictions exhibited systematic bias patterns that compromise drug repurposing applications. We compared prediction scores between drugs for which indications were represented in the training set versus drugs with no prior disease indications. Drugs with indications in the training set demonstrated a substantial positive bias, with approximately 70% of all high-confidence predictions originating from drugs seen during training (Figure 4A). Drugs not seen in the training set predominantly received lower prediction probabilities (0.4–0.5 range), whereas drugs seen in the training set showed more evenly distributed prediction scores across the full range. This training bias was accompanied by a bias towards more highly connected drugs (hubs) (Figure 4B). While most drugs received predictions for relatively few diseases, a small subset of highly-connected drugs seen in the training set were predicted as drug repurposing candidates for a large number diseases (up to 2000), as shown in (Figure S2). Thus, the model appears to overgeneralizes connectivity patterns present in the training graphs.

**Fig. 4.**
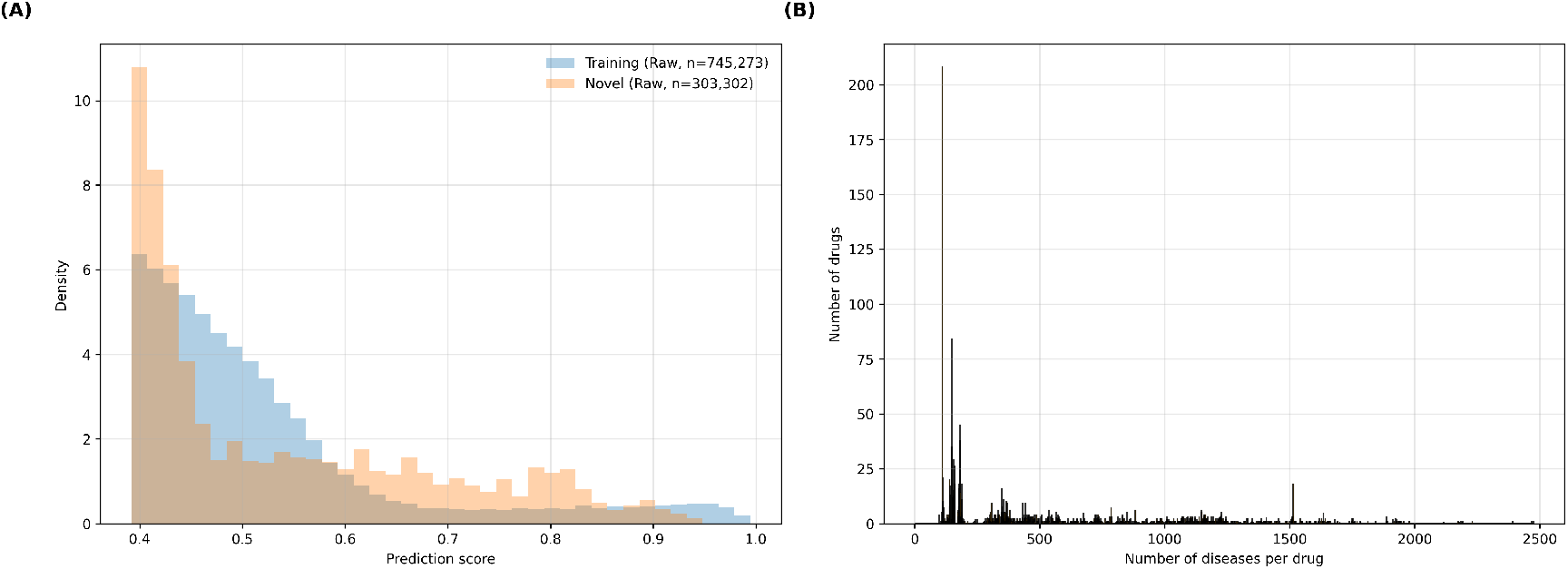
Bias diagnostics of model predictions from the TransformerConv model. (A) Distributions of model prediction scores, comparing links involving drugs present in the training set (blue) with those involving drugs absent from the training set (orange). (B) Frequency distribution of drug connectivity in predictions. X-axis shows the number of diseases per drug; Y-axis shows the count of drugs with that connectivity level.

Beyond these qualitative observations, we also obtained quantitative evidence from correlation analyses for systematic frequency bias in model predictions (Figure S3). Drug-disease pairs with higher combined frequencies in the training dataset (drug frequency × disease frequency) receive higher prediction scores (Spearman *ρ* = 0.61 (95% CI: [0.604, 0.611], p *<* 0.001); Pearson *r* = 0.32 (95% CI: [0.313, 0.322], p *<* 0.001); log-log Pearson *r* = 0.58 (95% CI: [0.579, 0.586], p *<* 0.001)). The larger rank-based and log-transformed correlations indicate a non-linear, heavy-tailed relationship between training frequency and prediction scores (Figure S3: (A-B) scatter plots, (C) distribution). However, correlations are still moderate, indicating that the model has learned beyond simple frequency patterns.

### Attribution Analysis

To evaluate the biological relevance of our predictions and gain insight into the reasoning of the model, we applied GNNExplainer to the TransformerConv results (Table 7). GNNExplainer analysis with different neighborhood sizes revealed significant differences in node type attribution patterns. 1-hop neighborhoods encompassed nodes with direct edges to either the target drug or disease, including immediate drug targets, drug type, disease-associated genes, and therapeutic areas. 2-hop analysis reveals indirect relationships by including intermediate biological entities, such as shared pathways and common targets that connect drugs and diseases through one additional step. For example:

**Table 7.** GNNExplainer-Based Interpretation of TransformerConv Predictions Across Neighborhood Sizes.

| Metric | 1-Hop Importance | 2-Hop Importance |
| --- | --- | --- |
| Node score range | [0.298–0.673] | [0.276–0.701] |
| *Node Type (95% CI):* |  |  |
| Drug | 0.406 [0.394–0.450] | 0.384 [0.383–0.385] |
| Disease | 0.435 [0.383–0.486] | 0.382 [0.382–0.382] |
| Gene | 0.383 [0.382–0.385] | 0.423 [0.418–0.429] |
| DrugType | 0.427 [0.373–0.505] | 0.484 [0.475–0.492] |
| TherapeuticArea | 0.378 [0.367–0.393] | 0.437 [0.430–0.445] |
| Pathway | – | 0.383 [0.382–0.383] |
| *Baseline Comparison:* |  |  |
| p-value | < 0.001 | < 0.001 |
| *Faithfulness Testing:* |  |  |
| Performance drop (attributed) | 0.204 ± 0.123 | 0.266 ± 0.170 |
| Performance drop (random) | 0.110 ± 0.119 | 0.110 ± 0.117 |
| p-value | < 0.001 | < 0.001 |

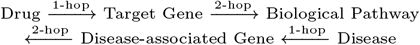

Gene importance increased from 0.383 to 0.423 in the expanded neighborhoods. Pathway nodes, absent in 1-hop attributions, emerged with moderate importance (0.383) in 2-hop analysis, representing the primary difference between the approaches. This indicates that genes become more predictive when considered within a pathway context rather than as isolated drug targets. The wider confidence intervals for 1-hop reflect higher variability due to smaller neighborhood size and fewer contextual nodes, whereas the narrow 2-hop neighborhoods provide more stable attribution patterns by incorporating additional biological context.

In our two-step validation framework, baseline comparison revealed that GNNExplainer produces systematically different attribution patterns compared to random baselines (Mann-Whitney U test, p *<* 0.001). Faithfulness testing demonstrated that removing top-attributed nodes caused significantly greater performance drops than removing random nodes (2-hop: 0.266 ± 0.170, 1-hop: 0.204 ± 0.123, Wilcoxon signed-rank test, p *<* 0.001). Together, these results indicate that GNNExplainer produces selective, non-uniform attributions where nodes receiving high importance scores are causally relevant to model predictions, demonstrating meaningful explanation quality rather than random noise.

## Discussion

We presented a unified evaluation framework for GNN-based drug repurposing, dubbed KG-Bench. KG-Bench enables fair, reproducible comparison of graph-based deep learning models for the prediction of drug repurposing candidates. Our approach ensures that all benchmark components adhere to the FAIR principles: datasets are assigned persistent identifiers and registered in public repositories for findability; open-source code and comprehensive documentation are accessible from our GitHub repository; interoperability is achieved through implementation of established biomedical ontologies and unified data schemas. To ensure reproducibility and consistency across experiments, we recommend that users leverage the cleaned and preprocessed dataset provided with our framework, which includes standardized training, validation, and test splits. This setup enables direct comparison with our reported results and facilitates fair benchmarking of new models. Alternatively, researchers may choose to follow our documented preprocessing pipeline to generate their own splits, allowing for flexibility in adapting the framework to novel datasets or experimental conditions. In both cases, the modular design of our framework supports integration of custom GNN architectures and evaluation metrics, making it suitable for a wide range of drug repurposing applications.

One of the most critical contributions of this work is addressing the pervasive data leakage problem in drug repurposing benchmarks. Data leakage often occurs when related entities, such as derivative drugs or disease terms that are hierarchically related, appear across training, validation, and test sets. This leads to overly optimistic performance estimates. By removing redundant drug variants, collapsing disease identifiers with a hierarchical relationship, and ensuring strict separation of edges between training, validation and test sets, we provide a benchmark that reflects realistic prediction scenarios.

The results in Table 6 show performance differences among the three GNN architectures, which can be partially attributed to their design principles, such as attention mechanisms and neighborhood sampling strategies. TransformerConv achieved the highest performance, reflecting the advantage of its attention mechanism in weighting informative neighbors in a heterogeneous KG. This capability is particularly important in a scale-free graph where connectivity varies widely, allowing TransformerConv to exploit both structural and feature diversity effectively. GraphSAGE performed slightly worse. It benefits from neighborhood sampling that improves generalization in sparse regions, but lacks the fine-grained weighting of TransformerConv. GCN showed the lowest performance, likely because its uniform aggregation dilutes signal from highly connected hubs, making it more reliant on topology alone. Despite architectural differences, the three models consistently identified similar novel drug-disease associations (Table S1).

Table 5 clearly shows the effects of different negative sampling ratios on model performance. As the ratio of negative to positive samples increased from 1:1 to 1:100, the APR dropped sharply across all models as expected. This decline highlights the sensitivity of precision to class imbalance and underscores that the choice of negative sampling strategy is a fundamental challenge in KG-based drug repurposing. Since KGs only contain known associations, it is unclear which unobserved drug–disease pairs represent true negatives and which represent pairs yet to be discovered. We chose random negative sampling in this benchmark framework for its simplicity, neutrality, and methodological consistency. Alternative strategies, such as structure-based, semantic distance-based, and ranking-based methods, have been proposed [39, 40, 41, 42], but these approaches can introduce systematic biases and may miss unexpected yet valid associations. Random sampling avoids such assumptions and ensures fair comparisons across models.

Bias is a prevalent issue in graph-based predictions of drug repurposing candidates. Models often perform better on well-studied diseases such as cancer or diabetes[43], and predictions for popular drugs with extensive literature tend to be more accurate[44]. This trend may be attributed to the higher representation frequency of these entities in the training data, which facilitates the learning of more robust embeddings and results in elevated model confidence scores. While new predictions may involve fewer connected entities or require the model to infer outside of the established hub patterns, resulting in systematically lower confidence scores. This challenge is closely related to the problem of one-shot prediction, which refers to the challenge of making accurate predictions for drugs or diseases that appear only once or very few times in the training data. Models like TxGNN[45] have addressed this kind of issue by incorporating disease similarity into the graph structure, enabling the model to generalize better to underrepresented diseases and improve prediction accuracy for novel drug–disease pairs. This bias highlights the need for bias-aware evaluation metrics. Detecting and reducing such biases is essential for ensuring fairness and generalisability in predictive models.

The types of nodes and the number of features in the benchmark KG are currently limited. Future benchmarks using the KG-Bench framework could include more features from Open Targets. Textual descriptions could be processed with language models like BioBERT[46] or Word2Vec[47] to generate text embeddings. Another promising extension to the benchmarking framework would be the implementation of edge weights, which assist the model in capturing the variable strengths of biological relationships.

To conclude, the KG-Bench framework implements a retrospective validation strategy for accurate benchmarking and provides a comprehensive performance assessment using multiple metrics, including AUC-ROC, precision–recall curves, and measures for bias and node attribution. Standardized and fair benchmarking addresses a critical gap in the field, where inconsistent methodologies and data leakage issues have complicated a fair comparison of published algorithms. Appropriate validation and benchmarking strategies will ultimately increase the proportion of drug repurposing candidates that are successful in clinical development[48, 49].

## Conclusion

Our contributions can be summarized as follows:

- **Open Targets Knowledge Graph:** We constructed a harmonized biomedical KG from Open Targets, integrating drugs, diseases, targets, and contextual annotations, and applied strict preprocessing to prevent data leakage from hierarchical ontology.
- **GNN Benchmarking:** We systematically evaluated multiple GNN architectures under consistent conditions, including interpretability analysis and ablation studies.
- **Framework Usability:** New users can either use the provided benchmark data for direct model comparison or reuse the preprocessing pipeline to generate custom datasets for their own models.

## Code availability

The code and data used in this study is available at https://github.com/cmbi/Benchmark_GNN_OpenTargets

---

**Fig. S1.**
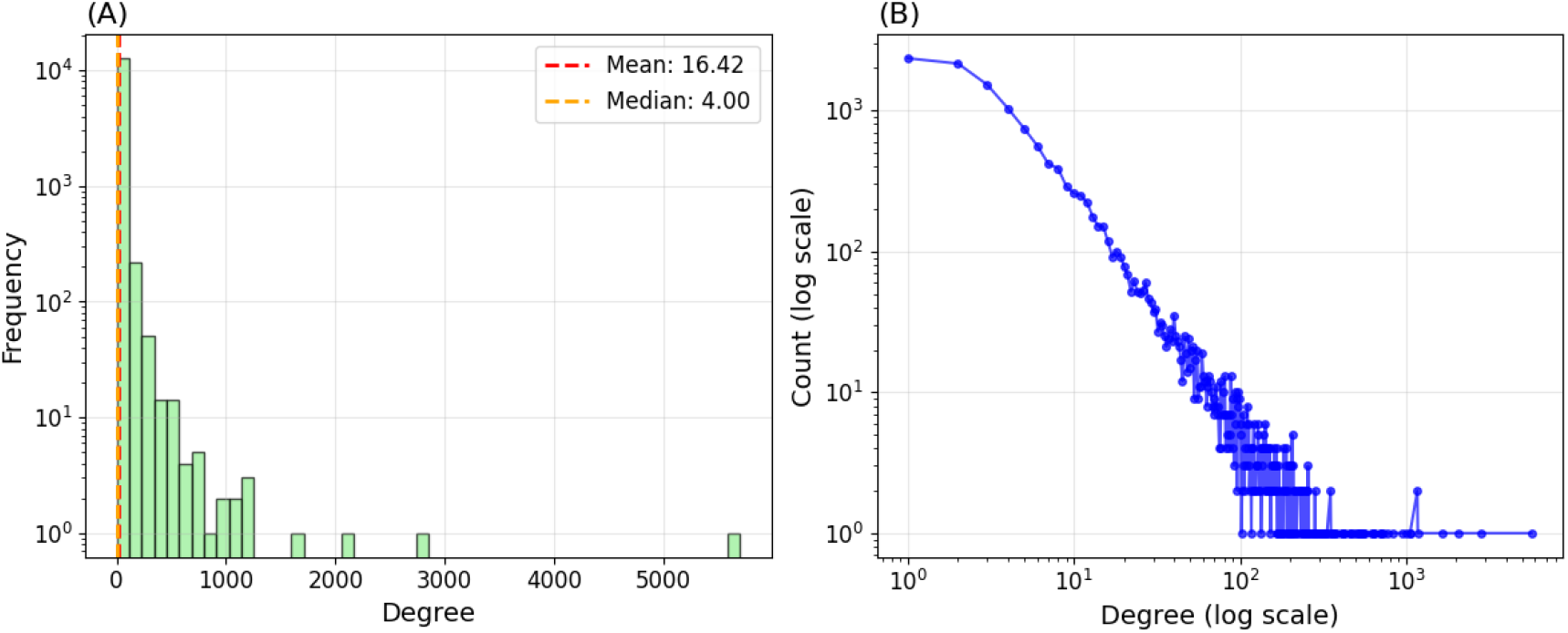
Graph topology of the Open Targets KG. **(A)** Degree frequency distribution (log scale) demonstrates large heterogeneity in node connectivity. **(B)** Power-law relationship in log-log space indicates a scale-free graph structure.

**Table S1.** Test Set Model Performance Comparison Across Different Positive:Negative Ratios.

| Ratio | Model | TP | FP | FN | TN |
| --- | --- | --- | --- | --- | --- |
| 1:1 | GCN | 71 | 31 | 55 | 95 |
|  | GraphSAGE | 95 | 29 | 31 | 97 |
|  | TransformerConv | 83 | 16 | 43 | 110 |
| 1:10 | GCN | 71 | 181 | 55 | 1079 |
|  | GraphSAGE | 95 | 257 | 31 | 1003 |
|  | TransformerConv | 83 | 118 | 43 | 1142 |
| 1:100 | GCN | 71 | 1623 | 55 | 10977 |
|  | GraphSAGE | 95 | 2363 | 31 | 10237 |
|  | TransformerConv | 83 | 992 | 43 | 11608 |

**Fig. S2.**
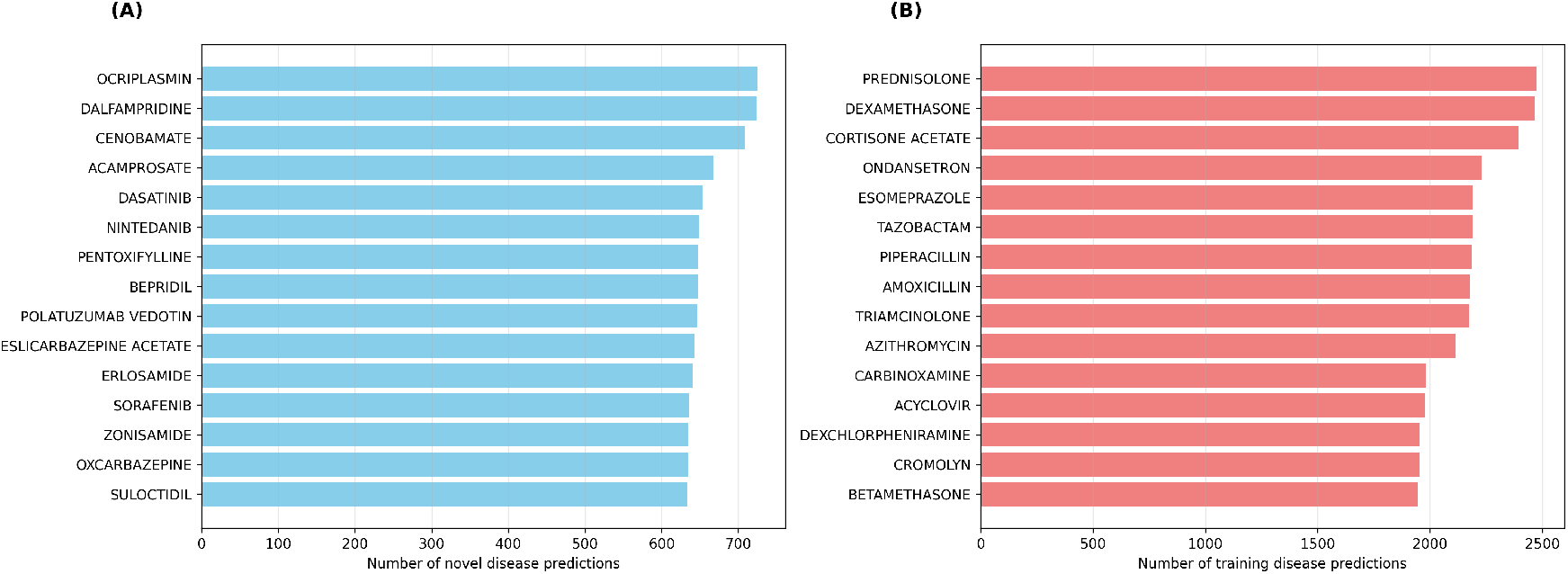
Analysis of model bias through drug-level prediction patterns. (A) Drugs absent from the training set ranked by the total number of disease predictions they involve. (B) Drugs present in the training set ranked by the total number of disease predictions they involve.

**Fig. S3.**
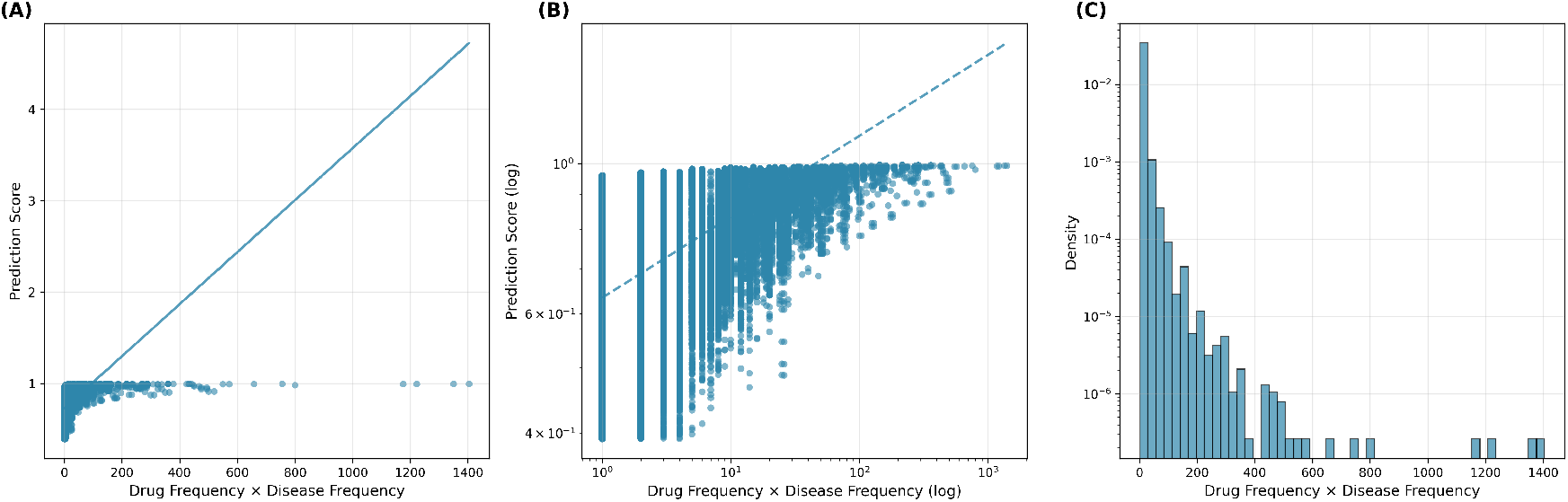
Correlation between model prediction scores and the combined training frequency of drug–disease pairs. (A) Scatter plot showing raw prediction scores versus the product of drug and disease frequencies in the training set. (B) Log–log scatter plot highlighting the heavy-tailed relationship between frequency and score. (C) Distribution of combined training frequencies across all evaluated pairs, illustrating the skewed connectivity pattern in the knowledge graph.

